# Epigenetic markers of adverse lifestyle identified among night shift workers

**DOI:** 10.1101/2022.07.13.499754

**Authors:** Paige M. Hulls, Daniel L. McCartney, Yanchun Bao, Rosie M. Walker, Frank de Vocht, Richard M. Martin, Caroline L. Relton, Kathryn L. Evans, Meena Kumari, Riccardo E. Marioni, Rebecca C. Richmond

## Abstract

**Background:** Epigenetic changes in the form of DNA methylation (DNAm) may act as biological markers of risk factors or adverse health states. We investigated associations between night shift work and established DNAm predictors of lifestyle, and compared them with those observed between night shift work and self-reported or conventionally-measured phenotypes.

**Methods:** In two cohort studies, Generation Scotland (GS) (n=7,028) and Understanding Society (UKHLS) (n=1,175), we evaluated associations between night shift work and four lifestyle factors (body mass index, smoking, alcohol, education) using both conventionally-measured phenotypes and DNA methylation-based scores proxying the phenotypes. DNA methylation-based measures of biological ageing were also generated using six established “epigenetic clocks”. Meta-analysis of GS and UKHLS results was conducted using inverse-variance weighted fixed effects.

**Results:** Night shift work was associated with higher BMI (0.79; 95%CI 0.02, 1.56; p=0.04) and lower education (-0.18; -0.30, -0.07; p=0.002). There was weak evidence of association between night shift work and DNAm scores for smoking (0.06, -0.03, 0.15; p=0.18) and education (-0.24; -0.49, 0.01; p=0.06) in fully adjusted models. Two of the epigenetic age measures demonstrated higher age acceleration among night shift workers (0.80; 0.42, 1.18; p<0.001 for GrimAge and 0.46; 0.00, 0.92; p=0.05 for PhenoAge).

**Conclusions:** Night shift work is associated with phenotypic and DNAm-based measures of lower education. Night shift work was also related to DNAm predictors of smoking and ageing.

## INTRODUCTION

Shift work has been referred to as *“a work activity scheduled outside standard daytime hours, where there may be a handover of duty from one individual or work group to another”*^1^. Typically, shift work has been associated with industries that require 24-hour operation, such as essential public services or for practical purposes. In recent years there has been an increase in the number of shift workers in other industries^1^ with approximately 19% of the working population engaged in shift work as their main job^2^ in the UK.

It has been argued that the introduction of shift work within wider industries does not consider the health and wellbeing costs to the individual shift worker^3^. Much of the research that investigates the health impacts of shift work has previously centred around circadian disruption, which can result in disturbed sleep and excessive sleepiness during the work shift^4^. However, adverse health behaviours have been found among shift workers which also put them at higher risk of disease ^5-7^ and shift work has been associated with higher risk of diseases^8-11^ than non-shift workers.

Recent studies have investigated epigenetic changes in the form of circulating DNA methylation (DNAm) as an objective measure for evaluating the potential health impact of shift work^12^. While these studies have typically focused on assessing individual sites in the genome (cytosine-phosphate-guanine “CpG” sites), recently DNAm scores derived from methylation levels at numerous CpG sites across the epigenome have been developed which can act as proxies for lifestyle exposures and may predict health outcomes^13^. Self-reported health behaviours (especially one-off measures) are subject to measurement error and bias ^14^, and using more objective measures of long-term exposure could identify more robust associations with shift work.

One group of DNAm scores aims to capture the epigenetic clock. DNA methylation age (DNAm Age) has been derived to provide an accurate estimate of biological age across a range of tissues, and at different life stages^15^. Estimated DNAm Age exceeding true chronological age is known as ‘age acceleration’ and studies suggest that DNAm age acceleration is associated with age-related health outcomes independent of chronological age^16^. As well as the clocks developed based on age, more recent ‘second generation’ epigenetic clocks have been developed based on lifestyle factors and biomarkers which have been found to be highly predictive of both health and lifespan^17,18^. Other DNAm scores which predict modifiable health, lifestyle and socio-economic factors include scores developed for alcohol consumption, smoking status, BMI and education^13^.

This study aimed to investigate associations of night shift work participation and a series of blood based DNAm predictors of ageing, BMI, smoking, alcohol and education within the *Generation Scotland* (GS) and *Understanding Society* (UKHLS) studies.

## MATERIALS AND METHODS

### 1.1. Generation Scotland (GS)

The Generation Scotland: Scottish Family Health Study is a prospective cohort study comprising participants from the general population across five regions of Scotland. The recruitment protocol and cohort characteristics are described in detail elsewhere^19^ and in the **Supplementary Methods**.

Blood DNAm was profiled using the Infinium MethylationEPIC BeadChip (Illumina Inc.) in two sample sets from GS: Set 1 and Set 2 (**Supplementary Methods**).

Methylation data were available for 777,193 CpGs measured in Set 1 (n=2,578 unrelated individuals) and 773,860 CpGs measured in Set 2 (n=4,450 unrelated individuals). The data sets had DNAm profiled at separate time points and quality control and normalisation was carried out separately. Details on the variables and how they were derived can be found in the **Supplementary Methods**.

### 1.2. Understanding Society (UKHLS)

The UK Household Longitudinal Study (UKHLS) (also known as Understanding Society) is a longitudinal panel survey of 40,000 UK households from England, Scotland, Wales and Northern Ireland. The recruitment protocol and cohort characteristics are described in detail elsewhere^20,21^ and in the **Supplementary Methods**.

Blood DNAm was profiled using the same Infinium MethylationEPIC BeadChip (Illumina Inc.) as used by GS. Methylation data was available for 837,487 CpGs in 1,175 individuals (**Supplementary Methods**). Details on the variables and how they were derived can be found in the **Supplementary Methods**.

### 1.3. DNA methylation scores

We derived four DNAm scores related to BMI, smoking, alcohol consumption and education which were based on CpG sites identified in a previous study^13^. Details of the DNAm scores are shown in Table 1. For each individual, DNAm scores were calculated as the sum of methylation values at each CpG multiplied by the effect sizes obtained in a previous study^13^.

**Table 1:** Origins of lifestyle DNA methylation scores employed in the current analysis.

| Phenotype | DNA methylation scores | Original publication | No. of CpG sites |
| --- | --- | --- | --- |
| Alcohol | Alcohol | Epigenetic prediction of complex traits and death<sup>13</sup> | 450 |
| Smoking | Smoking | Epigenetic prediction of complex traits and death<sup>13</sup> | 233 |
| BMI | BMI | Epigenetic prediction of complex traits and death<sup>13</sup> | 1109 |
| Education | Education | Epigenetic prediction of complex traits and death<sup>13</sup> | 373 |
| Ageing | AgeAccelHorvath | DNA methylation age of human tissues and cell types<sup>22</sup> | 353 |
|  | IEAA | DNA methylation age of human tissues and cell types<sup>22</sup> | 353 |
|  | AgeAccelHannum | Genome-wide methylation profiles reveal quantitative views of human ageing rates<sup>20</sup> | 71 |
|  | EEAA | DNA methylation-based measures of biological age: meta-analysis predicting time to death<sup>21</sup> | 71 |
|  | AgeAccelPheno | An epigenetic biomarker of ageing for lifespan and healthspan<sup>17</sup> | 513 |
|  | AgeAccelGrim | DNA methylation GrimAge strongly predicts lifespan and healthspan<sup>18</sup> | 1,030 |

CpG: cytosine-phosphate-guanine; IEAA: Intrinsic epigenetic age acceleration; EEAA: Extrinsic epigenetic age acceleration

We also derived six epigenetic biomarkers of ageing using previously published approaches^17,20-22^. (**Supplementary Methods**). In each case, age acceleration was defined as the residual obtained from regressing predicted age, as estimated by the epigenetic clock, on chronological age. This measure of age acceleration is independent of chronological age.

All methylation scores were standardized (mean = 0, standard deviation = 1) in both GS and UKHLS.

### 1.4. Statistical Analyses

We first assessed whether lifestyle factors were associated with night shift work. We performed linear regression of each phenotype (as the outcome variable) in relation to night shift work (as the exposure variable). This was with the exception for education, which was treated as the exposure variable given that education precedes night shift work. In GS, two models were run: Model 1 with adjustment for age and sex, and Model 2 with additional adjustment for other self-reported phenotypes (e.g., for smoking, the model was also adjusted for BMI, alcohol, and education). In UKHLS, two models were run: Model 1 with adjustment for age, sex, blood processing day and batch and Model 2 with additional adjustment for other self-reported phenotypes.

We subsequently assessed whether the lifestyle factors could be proxied with DNAm scores within GS and UKHLS. We performed linear regression of the phenotypes (exposure variable) and lifestyle DNAm scores for alcohol, smoking, education, and BMI (outcome variable). For these models, a logistic regression was performed. In GS, two models were run: Model 1 with adjustment for age, sex and 20 methylation principal components (PCs) and Model 2 with additional adjustment for other self-reported phenotypes (e.g., for smoking related DNAm scores, the model was also adjusted for BMI, alcohol, and education). In UKHLS, two models were run: Model 1 with adjustment for age, sex, blood processing day and batch and Model 2 with additional adjustment for other self-reported phenotypes.

We finally assessed whether night shift work was related to the lifestyle DNAm scores. We performed linear regression of each lifestyle methylation score (outcome) in relation to night shift work (exposure). This again was with the exception for education, which was treated as the exposure. In GS, three models were run: Model 1 with adjustment for age, sex and 20 methylation PCs; Model 2 had additional adjustments for other self-reported phenotypes (e.g., for smoking related DNAm scores, model adjusted for BMI, alcohol, and education); and Model 3 with further adjustment for the corresponding phenotype (e.g., for smoking related DNAm scores, the model was also adjusted for smoking). In UKHLS, three models were run: Model 1 with adjustment for age, sex, blood processing day and batch; Model 2 with additional adjustment of other self-reported; and Model 3 with further adjustment for the corresponding phenotype.

We assessed whether night shift work was related to any of the six epigenetic ageing measures. We performed linear regression of each epigenetic age acceleration (EEA) measure (as the outcome variable) in relation to night shift work.

In GS, two models were run: Model 1 with adjustment for sex and 20 methylation PCs and Model 2 with additional adjustment for smoking, BMI, education, and alcohol. In UKHLS, two models were run: Model 1 with adjustment for sex, blood processing day and batch and Model 2 with additional adjustment for smoking, BMI, education, and alcohol.

We conducted a series of meta-analyses of GS Sets 1 and 2 and UKHLS using an inverse-variance weighted fixed effects approach. For this, we used the binary measure of night shift work derived in GS and the current night shift work variable derived in UKHLS. The I^2^ statistic was used to assess heterogeneity across the studies.

We conducted meta-analyses of: i) the self-reported or conventionally measured (in the case of BMI) lifestyle factors in relation to night shift work, ii) the lifestyle DNAm scores in relation to night shift work and iii) the epigenetic age acceleration measures in relation to night shift work.

## RESULTS

### 2.1. Baseline characteristics

Summary characteristics of the participants from GS and UKHLS are presented in **Table 2**. 7,028 individuals with DNAm data were included from GS (n=2,578 in Set 1 and n=4,450 in Set 2) and 1,175 individuals from UKHLS. Set 1 and 2 of GS had a younger mean age (50.0±12.5 and 51.4±13.2) compared to UKHLS (58.0±15.0). There was a comparable balance of men and women across the three datasets: 38.6% males in GS Set 1, 43.7% in GS Set 2 and 41.6% in UKHLS. BMI was also broadly comparable: 27.4±5.5 kg/m^2^, 26.8±5.0 kg/m^2^ and 28.1±6.2 kg/m^2^, in GS Set 1, GS Set 2 and UKHLS, respectively. GS Set 1 had more current smokers whilst UKHLS had more former smokers and GS Set 2 had more never smokers. There was a higher proportion of daily drinkers in UKHLS (16.0%) compared with GS (12.4% in Set 1 and 13.2% in Set 2). There was also a higher proportion of less than monthly drinkers in UKHLS (25.2%) compared with GS (17.2% in Set 1 and 16.2% in Set 2). Years of full-time education was comparable between Set 1 and 2 of GS (13.6±3.4 and 13.8±3.4, respectively) whilst in UKHLS it was slightly lower (12.3±5.1). In UKHLS, 1.6% of participants (n=18) were currently working night shifts while 8.8% (n=103) had reported working night shifts over the previous 11 years. In GS Set 1, 8.1% (n=127) of participants reported working at night for >20 hours per week at the time of sampling, compared with 7.9% (n=193) in GS Set 2.

**Table 2:** Baseline characteristics of the participants in Generation Scotland and Understanding Society.

| Variables | Categories | Generation Scotland |  |  |  |  | Understanding Society |  |
| --- | --- | --- | --- | --- | --- | --- | --- | --- |
|  |  | Set 1<br>(n=2578) |  | Set 2<br>(n=4450) |  |  | Overall<br>(n=1175) |  |
|  |  | Mean | SD | Mean | SD |  | Mean | SD |
| **Age (years)** |  | 50.02 | 12.5 | 51.39 | 13.2 |  | 57.97 | 15.0 |
| **Body mass index (kg/m<sup>2</sup>)** |  | 27.24 | 5.4 | 26.76 | 4.9 |  | 28.11 | 6.2 |
| **Education (years of full-time education)** |  | 13.62 | 3.4 | 13.75 | 3.4 |  | 12.31 | 5.1 |
|  |  | N | % | N | % |  | N | % |
| **Sex** | *Male* | 1583 | 38.6 | 1944 | 43.7 |  | 489 | 41.6 |
|  | *Female* | 995 | 61.4 | 2506 | 56.3 |  | 686 | 58.4 |
| **Smoking status** | *Current* | 460 | 18.3 | 675 | 15.5 |  | 186 | 15.9 |
|  | *Former* | 771 | 30.7 | 1398 | 32.1 |  | 487 | 41.6 |
|  | *Never* | 1276 | 50.9 | 2276 | 52.3 |  | 498 | 42.5 |
| **Alcohol** | *Daily* | 167 | 12.4 | 334 | 13.2 |  | 164 | 16.0 |
|  | *More than weekly* | 728 | 54 | 1402 | 55.6 |  | 475 | 43.6 |
|  | *More than monthly* | 222 | 16.5 | 386 | 15.2 |  | 166 | 15.2 |
|  | *Less than monthly* | 231 | 17.2 | 411 | 16.2 |  | 275 | 25.2 |
| **Night shift work** | *0 hours per week* | 1015 | 64.9 | 1592 | 65.6 | *Never* | 1072 | 91.2 |
|  | *1-19 hours per week* | 422 | 27.0 | 643 | 26.5 | *Ever* | 103 | 8.8 |
|  | *>20 hours per week* | 127 | 8.1 | 193 | 7.9 | *Previous* | 79 | 6.9 |
|  |  |  |  |  |  | *Current* | 18 | 1.6 |

### 2.2. Lifestyle factors

#### 2.2.1. How does night shift work relate to conventionally measured lifestyle factors?

In GS, night shift work was inversely associated with alcohol frequency in both GS Set 1 and GS Set 2 (Model 1: Effect= -0.056; 95% CI -0.095, -0.017; p=0.006 category change per hour for GS Set 1, and - 0.039; -0.072, -0.006; p=0.019 for GS Set 2), although associations attenuated with adjustment for the other self-reported phenotypes in GS Set 2 (Model 2: -0.024; -0.057, 0.009; p=0.151) (**Table S1**). Night shift work was positively associated with smoking status in GS Set 1 (Model 1: 0.024; 0.006, 0.042; p=0.009 category change per hour) but were more weakly associated in GS Set 2 (Model 2: 0.012; -0.006, 0.030; p=0.216), and when adjusted for the other phenotypes (**Table S1**). There were positive associations between night shift work and BMI in both GS Set 1 (Model 1: 0.15; 0.03, 0.28; p=0.02 kg/m^2^ per hour) and GS Set 2 (Model 2: 0.23; 0.14, 0.32; p=1x10^-6^), although these associations attenuated when adjusted for the other phenotypes. There was an inverse association between night shift work and education in GS Set 2 (Model 1: -0.07; -0.13, -0.02; p=0.01 hour per year) which was less apparent in GS Set 1 (Model 1: -0.05; -0.12, 0.03; p=0.21), although there was stronger evidence of association of the binary night shift work variable with education, based on in both datasets (**Table S1**).

In UKHLS, no strong associations were found between current, ever, or previous night shift work and any of the lifestyle phenotypes, although effect estimates were typically in the same direction as in GS (**Table S2**).

In the GS-UKHLS combined meta-analysis there was evidence of a positive association between night shift work and BMI (0.79; 0.02, 1.56; p=0.04 kg/m^2^ difference between those who did and did not work night shifts) and an inverse association between night shift work and education (-0.18; -0.30, - 0.07; p=0.002 log odds per year of education) in the fully adjusted models (Model 2) (**Figure 1**). There was little evidence for an association between night shift work and either alcohol intake or smoking status in a meta-analysis across the three datasets (0.00; -0.19, 0.20; p=0.97 and 0.04; - 0.19, 0.27; p=0.73 category change between those who did and did not work night shifts, respectively). There was little evidence for heterogeneity between the study estimates (I^2^ < 5%). Results of the minimally adjusted models (Model 1) are shown in **Figure S1**).

**Figure 1:**
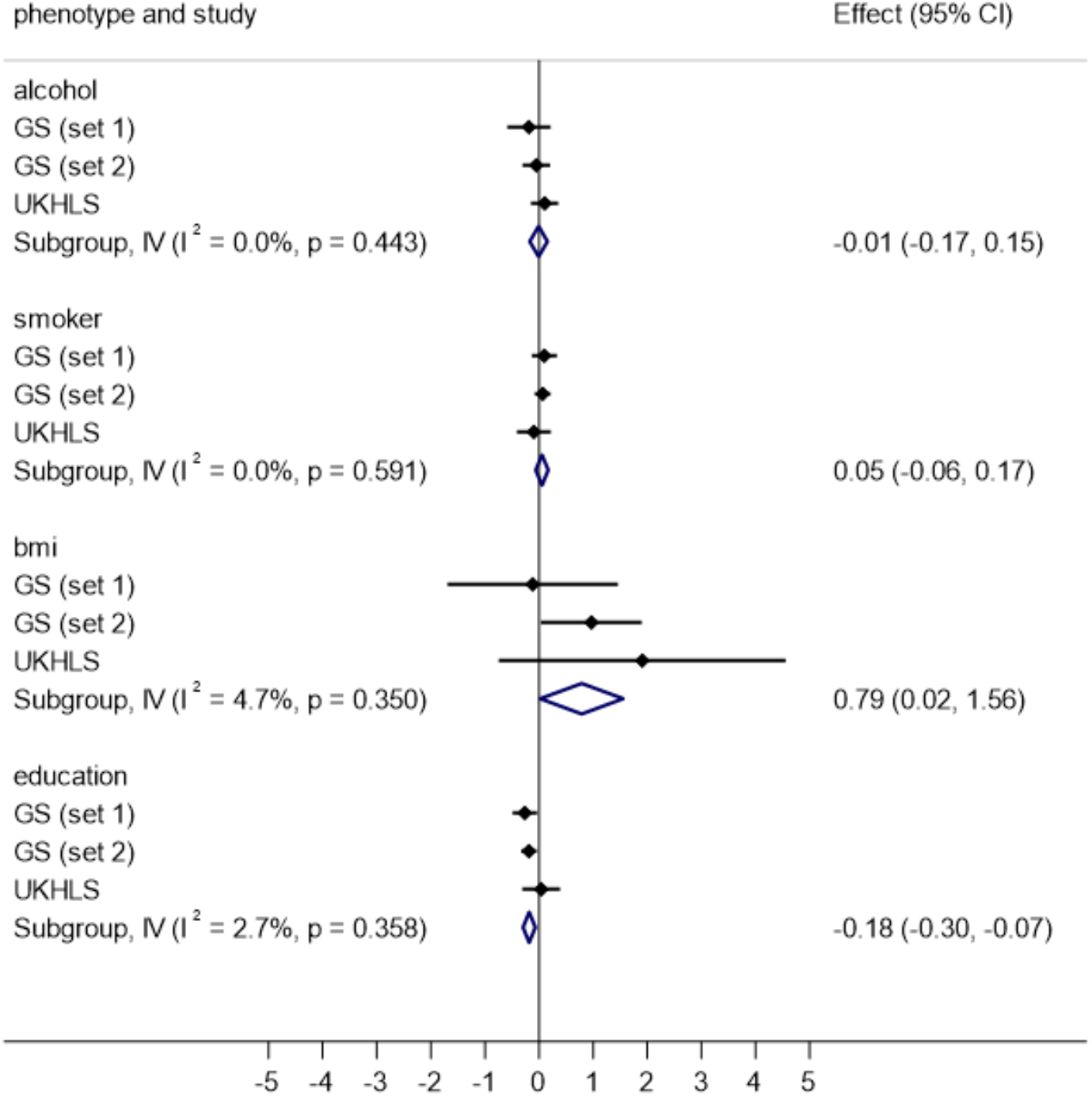
Associations between night shift work and phenotypes in Generation Scotland (GS) and Understanding Society (UKHLS) *. **Model 2: adjusted for age, sex, other self-reported phenotypes (e.g., for smoking, model adjusted for alcohol, body mass index and education)** For these models, education was treated as the exposure and shift work was the outcome; effect estimates are log odds ratios

#### 2.2.2. How are DNA methylation biomarkers associated with lifestyle factors in GS and UKHLS?

There were positive associations between each DNAm score and its respective lifestyle phenotype in both GS and UKHLS (**Table S3** and **S4**, respectively).

#### 2.2.3. Is night shift work associated with DNA methylation biomarkers?

In GS, there were no clear associations between night shift work and the alcohol DNAm score (**Table S5**). Night hours were positively associated with the smoking DNAm score in both GS Set 1 and GS Set 2 (Model 1: 0.04 SD; 0.02, 0.06; p=1.28x10^-4^ and 0.02 SD; 0.00, 0.04; p=0.01), although associations attenuated with further covariate adjustment in Models 2 and 3. Number of night hours was also positively associated with the BMI DNAm score in both GS Set 1 and GS Set 2 (Model 1: 0.03 SD; 0.01, 0.05; p=0.01 and 0.03 SD; 0.01, 0.04; p=0.04). Associations attenuated with further covariate adjustment in Models 2 and 3 (**Table S5**). Number of night hours was inversely associated with the education DNAm score in GS Set 1 (Model 1: -0.20 hours; -0.33, -0.08; p=0.001) and more weakly in GS Set 2 (Model 1: -0.09 hours; -0.19, 0.01; p=0.08). With further covariate adjustment, the association persisted in GS Set 1 but was attenuated in GS Set 2 (**Table S5**).

None of the night shift work measures were strongly associated with the lifestyle DNAm scores when UKHLS was assessed independently (**Table S6**). When estimates from GS were combined in a meta-analysis with UKHLS, there was little evidence of night shift work associations with the alcohol and BMI DNAm scores in the fully adjusted models (Model 3) (**Figure 2**). However, there was some evidence of an inverse association between night shift work and the education DNAm score in all three datasets (-0.24; -0.49, 0.01; p=0.06 log odds per year) (**Figure 2**). There was also weak evidence for a positive association between night shift work and the smoking DNAm score (0.06 SD, -0.03, 0.15; p=0.18). There was little evidence for heterogeneity between the study estimates in the meta-analysis (I^2^ = 0%). Results of the minimally adjusted models (Models 1 and 2) are shown in **Figures S2** and **S3**.

**Figure 2:**
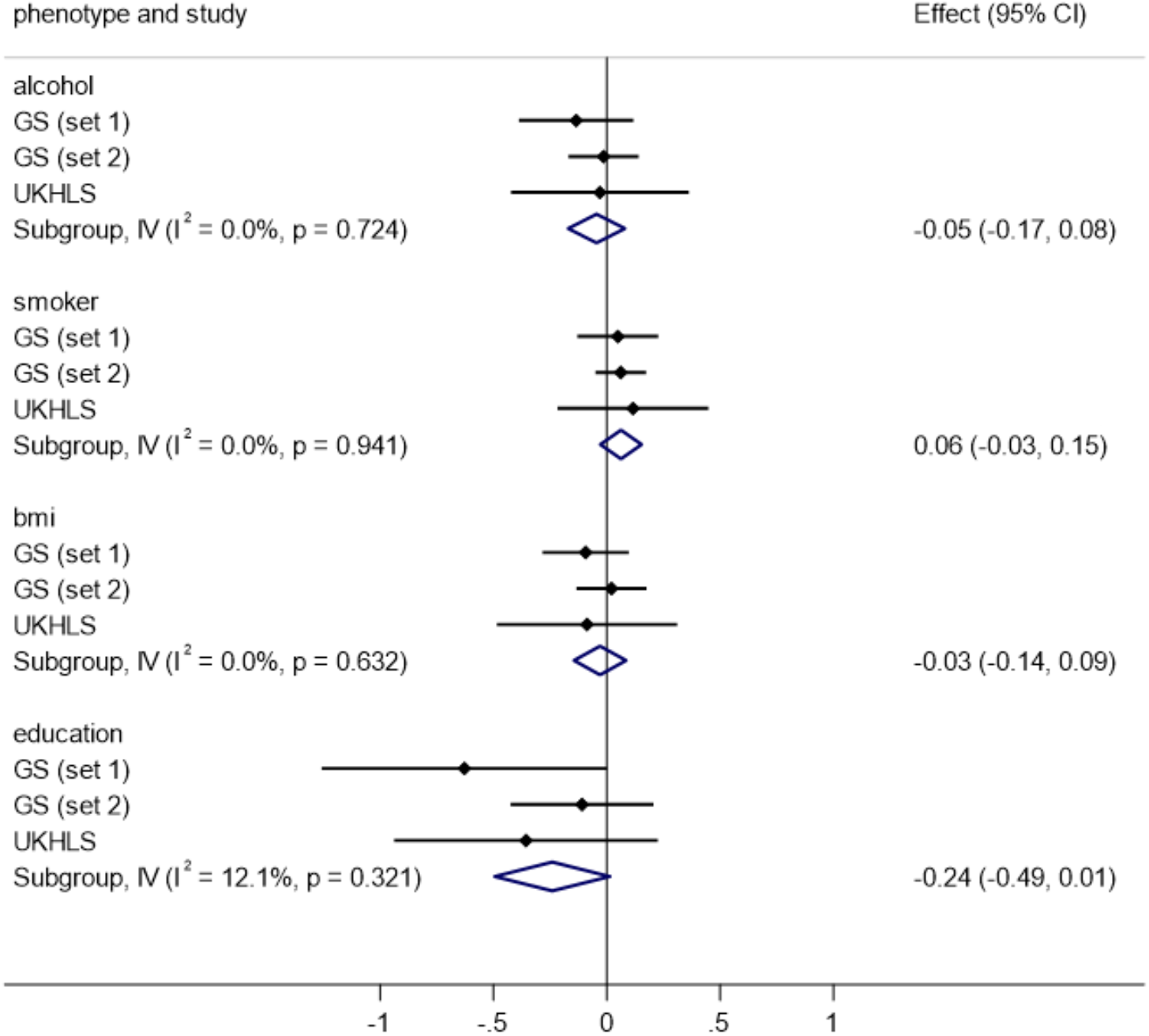
Associations between night shift work and methylation scores in Generation Scotland (GS) and Understanding Society (UKHLS) **\*Model 3: adjusted for age, sex, blood processing day, rack barcode, corresponding other self-reported phenotypes (e.g., for smoking DNAm, model adjusted for smoking, alcohol, body mass index and education)** **For these models, education was treated as the exposure and shift work was the outcome; effect estimates are log odds ratios**

### 2.3. Epigenetic ageing

There was evidence of an association between night shift work and GrimAge in GS, but not for the other epigenetic clocks. For GrimAge, the number of night hours was associated with higher age acceleration in both GS Set 1 (Model 1: 0.19 years, 0.11, 0.27; p=1.18x10^-5^) and GS Set 2 (Model 1: 0.10 years, 0.04, 0.16; p=9.45x10^-4^) **(Table S7)**. Associations with GrimAge acceleration remained, although were partially attenuated on adjustment for smoking, BMI, education and alcohol (***Table S7***). None of the night shift work measures were strongly associated with epigenetic age acceleration in UKHLS **(Table S8)**.

In the GS-UKHLS meta-analysis, night shift work was associated with a 0.80 year (0.42, 1.18; p<0.001) increase in GrimAge acceleration (**Figure 3**). There was also weak evidence of association with PhenoAge acceleration (0.46 years; 0.00, 0.92; p=0.05). The other four epigenetic clocks showed limited evidence of association. There was low heterogeneity between the study estimates in the meta-analysis (I^2^ < 50%).

**Figure 3:**
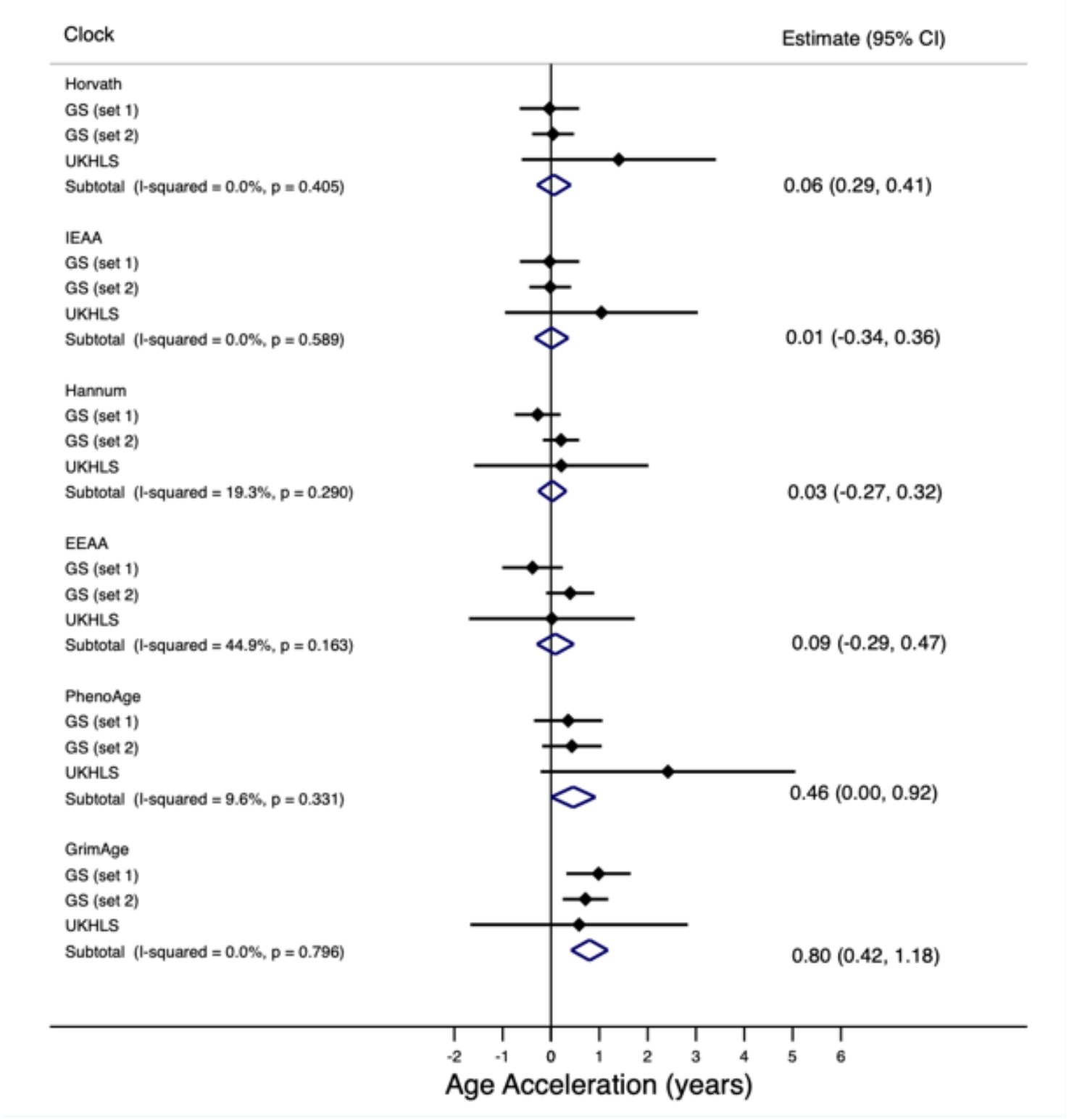
Associations between night shift work and epigenetic age acceleration in Generation Scotland (GS)and Understanding Society (UKHLS) **Model 2: Adjusted for sex, 20 methylation PCs, smoking, alcohol, body mass index and education** For these models, education was treated as the exposure and shift work was the outcome; effect estimates are log odds ratios

## DISCUSSION

We conducted analyses to investigate associations between night shift work and both phenotypic and DNAm markers in two cohorts. When we assessed phenotypic traits, we found that night shift work was associated with higher BMI and lower education. When assessing DNAm predictors of the same traits, there was similar evidence of association of night shift work with BMI and education DNAm scores. While the association with the BMI score was attenuated after adjusting for the corresponding phenotype, night shift work was nominally associated with education and smoking in fully adjusted models. Furthermore, two of the epigenetic age measures, GrimAge and PhenoAge, demonstrated higher age acceleration among night shift workers.

The observational associations of night shift work with lower education and higher BMI have been previously reported with comparable effect sizes^23^. While we did not find evidence of a phenotypic association with reported smoking, the association between night shift work and the smoking methylation score is consistent with previous studies reporting that smoking behaviour is more common with shift work comparison to day work^8,24-27^. One study also found that shift workers were also more likely to start smoking in comparison to their counterpart day workers^28^. We also found little evidence of an association between night shift work and either self-reported or methylation measures of alcohol; however, previous research has found that shift workers were more likely to drink heavily^7^.

There is growing evidence to suggest that DNAm-based measures are useful for health and lifestyle profiling^29^. Furthermore, several studies have shown that methylation predictors can provide a more accurate measurement of exposure than those based on self-report^30^. For example, previous studies have shown that smoking methylation scores may provide a more accurate measure of true exposure compared with self-reported smoking^18,30^, possibly due to erroneous self-reporting, the broad categories for reporting exposure, or because DNAm is able to capture long-term biological changes as a result of smoking as well as secondary smoking. This is supported in part by our findings that the education and smoking scores were still weakly associated with night shift work even after adjusting for the corresponding self-reported exposures.

Night shift work was also associated with higher epigenetic age acceleration, as measured by two epigenetic clocks: one predictive of mortality (GrimAge)^18^ and the other predictive of physiological dysregulation (PhenoAge)^17^. White *et al* (2019)^12^ also found associations between the length of time working night shifts and increased PhenoAge^31^, although GrimAge was not investigated. Circadian oscillators have been found to contribute to epigenetic ageing^32^ and there is emerging evidence that DNAm age estimators relate to circadian rhythm^33^. However, is should be noted that we did not find that night shift work was associated with the Hannum and Horvath clocks, which were designed to estimate chronological age^20,22^. This absence of association is in accordance with findings from White *et al*^31^ and suggests that intrinsic circadian processes are unlikely to underlie the associations observed^17,18^.

### Strengths and limitations

One of the main strengths of the study is the use of GS, an epidemiological cohort study with a large sample size with DNAm data which has also captured information on night shift work. Meta-analysing associations with those from the UKHLS dataset also improved power to detect associations between night shift work, lifestyle factors and DNAm predictors and indicated consistency in associations across studies. Furthermore, we have the use of both self-reported and DNAm markers for the same exposure, so were able to directly compare.

There were some limitations to the study. The DNAm scores used in this study were developed in GS Set 1, which might lead to overfitting of the models. However, by using the second set of participants in GS and the independent UKHLS datasets, we hope that this should minimise any potential overfitting issues. There was also limited evidence for heterogeneity in the associations observed between GS set 1 and the other studies.

We specifically assessed night shift work rather than other forms of shift work (such as morning/evening only or rotating shift work) in relation to DNA methylation. This is because of the previous evidence suggesting that night shift work is likely to be particularly disruptive to biological processes and to have implications for adverse health. Whilst rotating shift work has also been linked with circadian disruption, we did not specifically investigate this group of workers. There is limited information on the intensity of shift work per night, e.g. whether the participant works 7pm-7am three days a week or 7pm-10pm every day, or the direction of shift rotation (forward or backward rotating) which may have a differential biological impact^34^.

We are unable to make conclusions regarding causality of the associations observed. We cannot exclude reverse causation since the analysis in GS and that based on current night shift work in UKHLS was assessed cross-sectionally. Given education preceeds shift work status, the association implies that those with less education engage in occupations that include night shift work. This is in line with a previous study which found an inverse association between a genetic (rather than epigenetic) risk score for higher education and shift work participation^35^.

To support our findings, similar analysis should be performed in larger cohorts with DNAm data. Future studies should also look at evaluating whether these biomarkers could provide insights into the potential effects of night shift work on subsequent health outcomes, e.g., cardiometabolic diseases and cancer.

## CONCLUSIONS

In over 8,000 participants from two cohort studies, night shift work was associated with both phenotypic and DNA methylation-based measures of higher BMI and lower education. DNAm predictors of smoking and ageing were also related to night shift work. Epigenetic measures may provide insights into the health and lifestyle profiles of night shift workers.

## SUPPLEMENTARY MATERIALS

Supplementary Table 1: Associations between night shift work and phenotypes in Generation Scotland

Supplementary Table 2: Associations between night shift work and phenotypes in Understanding Society

Supplementary Table 3: Associations between phenotypes and methylation scores in Generation Scotland

Supplementary Table 4: Associations between phenotypes and methylation scores in Understanding Society

Supplementary Table 5: Associations between night shift work and methylation scores in Generation Scotland

Supplementary Table 6: Associations between night shift work and methylation scores in Understanding Society

Supplementary Table 7: Associations between night shift work and measures of epigenetic age acceleration in Generation Scotland

Supplementary Figure 1: Associations between night shift work and phenotypes in Understanding Society and Generation Scotland – Model 1

Supplementary Figure 2: Associations between night shift work and methylation scores in Understanding Society and Generation Scotland – Model 1

Supplementary Figure 3: Associations between night shift work and methylation scores in Understanding Society and Generation Scotland – Model 2

## AUTHOR CONTRIBUTIONS

Conceptualization, Rebecca Richmond; Data curation, Daniel L. McCartney and Yanchun Bao; Formal analysis, Paige Hulls, Daniel L. McCartney and Rebecca Richmond; Supervision, Frank De Vocht and Richard Martin; Visualization, Richard Martin; Writing – original draft, Paige Hulls and Rebecca Richmond; Writing – review & editing, Paige Hulls, Daniel L. McCartney, Yanchun Bao, Rosie Walker, Frank De Vocht, Caroline Relton, Kathryn Evans, Meena Kumari, Riccardo Marioni and Rebecca Richmond.

## INSTITUTIONAL REVIEW BOARD STATEMENT

Data governance was provided by the METADAC data access committee, funded by ESRC, Wellcome, and MRC (2015–2018: Grant Number MR/N01104X/1 2018–2020: Grant Number ES/S008349/1).

Ethical approval for Understanding Society was obtained from the National Research Service (Understanding Society – UK Household Longitudinal Study: A Biosocial Component, Oxfordshire A REC, Reference: 10/H0604/2). All our consents can be found here: https://www.understandingsociety.ac.uk/documentation/health-assessment/fieldwork-documents.

## INFORMED CONSENT STATEMENT

Informed consent was obtained from all subjects involved in the study.

## DATA AVAILABILITY STATEMENT

According to the terms of consent for GS participants, access to data must be reviewed by the GS Access Committee. Applications should be made to.

