## Supplementary Tables for "Epigenetic markers of adverse lifestyle identified among night shift workers"

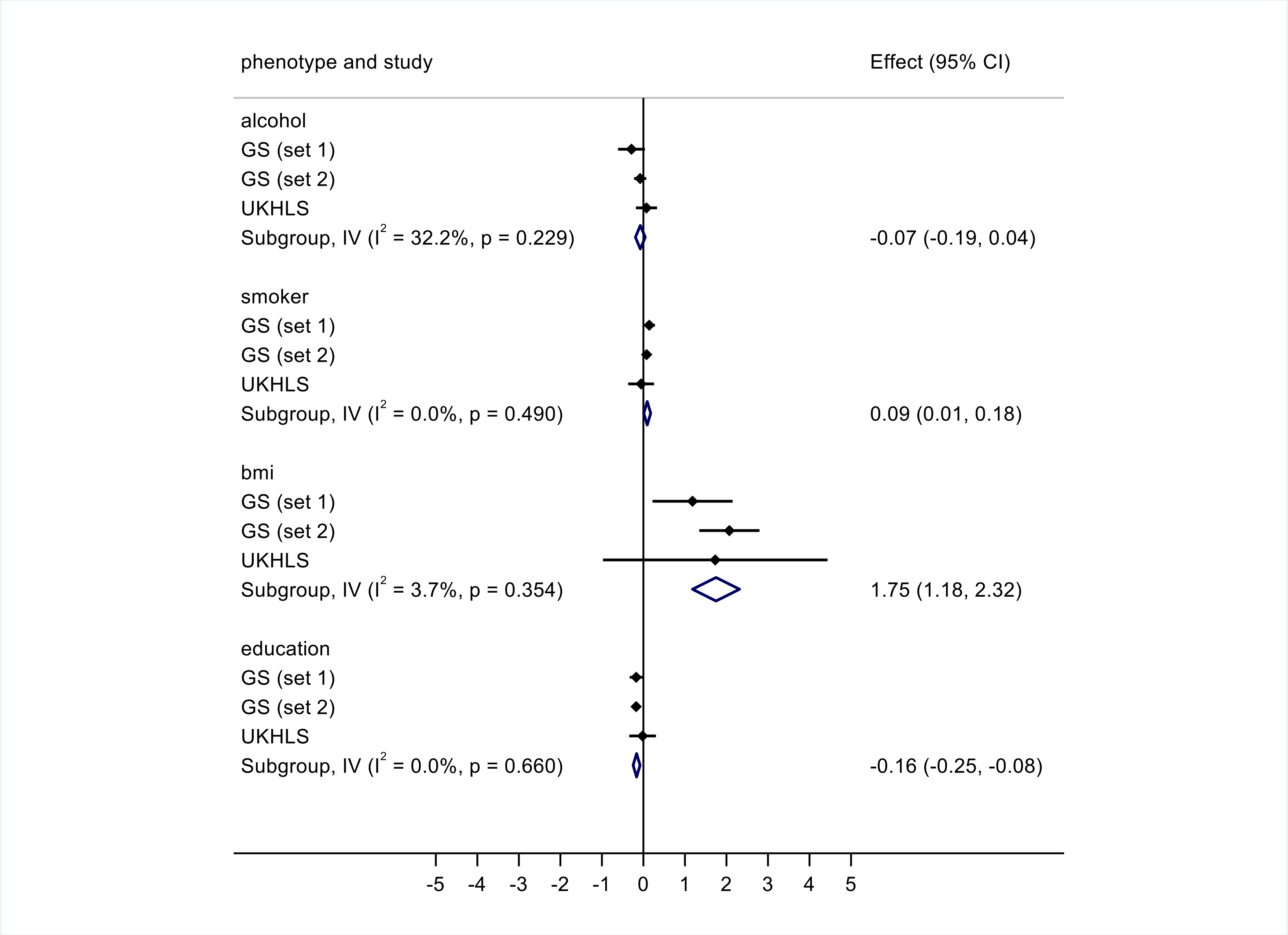


Supplementary Figure 1: Associations between night shift work and phenotypes in Understanding Society and Generation Scotland – Model 1

In these models, alcohol, smoking and BMI were treated as the outcome and shift work as the exposure (effect estimates are beta values) while education was treated as the exposure and shift work as the outcome (effect estimates are log odds ratios)

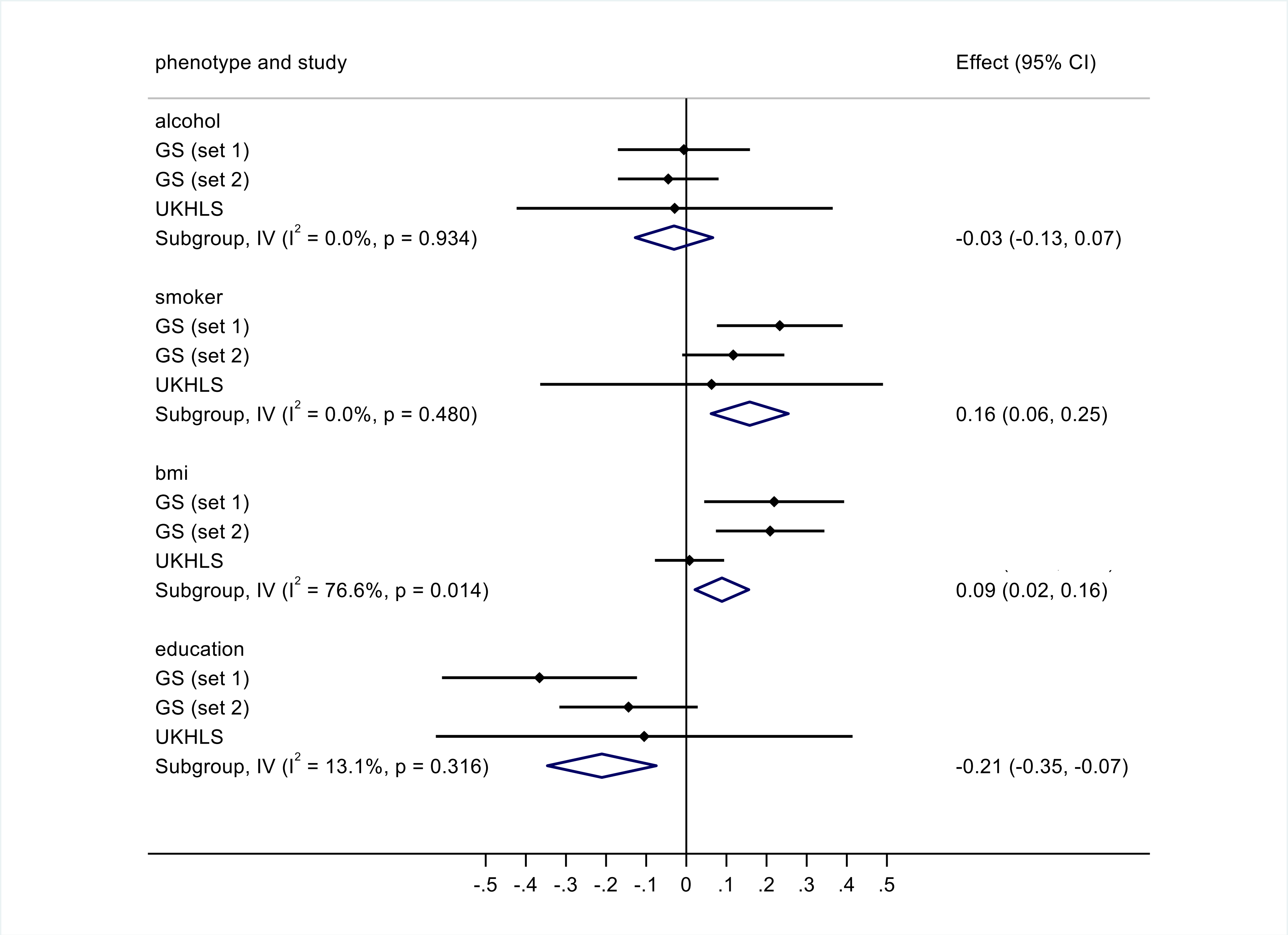


Supplementary Figure 2: Associations between night shift work and methylation scores in Understanding Society and Generation Scotland – Model 1

In these models, alcohol, smoking and BMI were treated as the outcome and shift work as the exposure (effect estimates are beta values) while education was treated as the exposure and shift work as the outcome (effect estimates are log odds ratios)

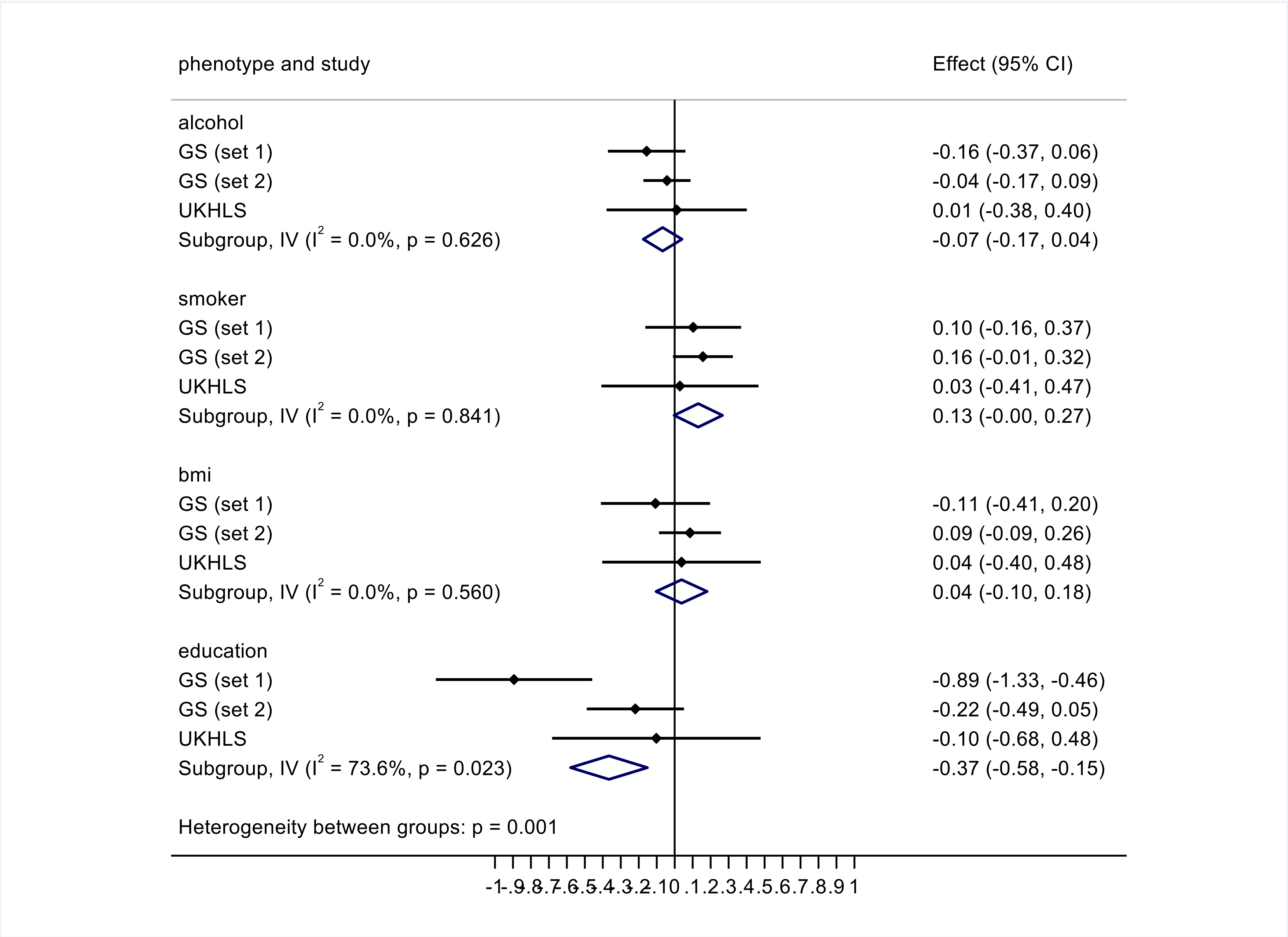


Supplementary Figure 3: Associations between night shift work and methylation scores in Understanding Society and Generation Scotland – Model 2

In these models, alcohol, smoking and BMI were treated as the outcome and shift work as the exposure (effect estimates are beta values) while education was treated as the exposure and shift work as the outcome (effect estimates are log odds ratios)
