## Supplementary Figures for "Epigenetic markers of adverse lifestyle identified among night shift workers"

Supplementary Table 1: Associations between night shift work and phenotypes in Generation Sc

| | | Set 1 (V | | | |
|---|---|---|---|---|---|
| Shiftwork | Phenotype | Model 1* (n=792-1563) | | | |
| | | effect | se | p-value | 95% CI |
| Night hours | alcohol | -0.056 | 0.020 | 0.006 | -0.095 |
| | smoker | 0.024 | 0.009 | 0.009 | 0.006 |
| | bmi | 0.152 | 0.063 | 0.016 | 0.028 |
| | education° | -0.048 | 0.039 | 0.214 | -0.124 |
| Binary night shift work | alcohol | -0.287 | 0.165 | 0.082 | -0.610 |
| | smoker | 0.142 | 0.070 | 0.044 | 0.005 |
| | bmi | 1.184 | 0.493 | 0.016 | 0.218 |
| | education°° | -0.174 | 0.080 | 0.029 | -0.331 |

*covariates: age and sex

**covariates: age, sex, other self-reported phenotypes (e.g. for smoking, model adjusted for alc

°For these models, education was treated as the exposure and shift work was the outcome; °°e

:otland

| Vave 1) | | | | | | |
|---|---|---|---|---|---|---|
| | Model 2** (n=480-620) | | | | | |
| | effect | se | p-value | 95% CI | | effect |
| -0.017 | -0.053 | 0.025 | 0.035 | -0.102 | -0.004 | -0.039 |
| 0.042 | 0.020 | 0.015 | 0.172 | -0.009 | 0.049 | 0.013 |
| 0.276 | 0.001 | 0.105 | 0.991 | -0.205 | 0.208 | 0.227 |
| 0.028 | -0.070 | 0.051 | 0.174 | -0.170 | 0.030 | -0.073 |
| 0.036 | -0.188 | 0.204 | 0.356 | -0.588 | 0.212 | -0.079 |
| 0.279 | 0.098 | 0.118 | 0.406 | -0.133 | 0.329 | 0.078 |
| 2.149 | -0.119 | 0.804 | 0.883 | -1.694 | 1.457 | 2.071 |
| -0.017 | -0.265 | 0.116 | 0.022 | -0.492 | -0.038 | -0.175 |

:ohol, body mass index and education)
ffect estimates are log odds ratios

| Set 2 (Wave 3) | | | | | | |
|---|---|---|---|---|---|---|
| Model 1* (n=1142-2415) | | | | Model 2** (n=1104- | | |
| se | p-value | 95% CI | | effect | se | p-value |
| 0.017 | 0.019 | -0.072 | -0.006 | -0.024 | 0.017 | 0.151 |
| 0.007 | 0.070 | -0.001 | 0.027 | 0.012 | 0.009 | 0.216 |
| 0.046 | 1.07E-06 | 0.136 | 0.318 | 0.100 | 0.061 | 0.101 |
| 0.026 | 0.006 | -0.125 | -0.022 | -0.074 | 0.035 | 0.035 |
| 0.075 | 0.289 | -0.226 | 0.068 | -0.051 | 0.130 | 0.695 |
| 0.057 | 0.168 | -0.034 | 0.190 | 0.068 | 0.075 | 0.366 |
| 0.371 | 2.64E-08 | 1.344 | 2.797 | 0.968 | 0.476 | 0.042 |
| 0.053 | 9.87E-04 | -0.279 | -0.071 | -0.188 | 0.072 | 0.009 |

| | |
|---|---|
| -1400) | |
| 95% CI | |
| -0.057 | 0.009 |
| -0.006 | 0.030 |
| -0.019 | 0.220 |
| -0.143 | -0.005 |
| -0.306 | 0.204 |
| -0.079 | 0.215 |
| 0.035 | 1.900 |
| -0.329 | -0.047 |

Supplementary Table 2: Associations between night shift work and phenotypes in Understandi

| | CURRENT NIGHT SHIFT | | | | | |
|---|---|---|---|---|---|---|
| Phenotype | Model 1 (n=1,086-1,167)* | | | | | |
| | effect | se | p-value | 95% CI | | effect |
| alcohol | 0.073 | 0.13 | 0.574 | -0.182 | 0.329 | 0.103 |
| smoker | -0.055 | 0.159 | 0.725 | -0.367 | 0.256 | -0.097 |
| bmi | 1.73 | 1.38 | 0.212 | -0.986 | 4.44 | 1.908 |
| education° | -0.02 | 0.164 | 0.901 | -0.342 | 0.302 | 0.04 |

| | EVER NIGHT SHIFT | | | | | |
|---|---|---|---|---|---|---|
| Phenotype | Model 1 (n=1,086-1,167)* | | | | | |
| | effect | se | p-value | 95% CI | | effect |
| alcohol | -0.006 | 0.066 | 0.924 | -0.137 | 0.124 | 0.009 |
| smoker | 0.075 | 0.336 | 0.336 | -0.079 | 0.23 | 0.042 |
| bmi | 0.961 | 0.695 | 0.167 | -0.403 | 2.32 | 0.943 |
| education° | -0.005 | 0.073 | 0.94 | -0.148 | 0.137 | -0.0004 |

| | PREVIOUS NIGHT SHIFT | | | | | |
|---|---|---|---|---|---|---|
| Phenotype | Model 1* | | | | | |
| | effect | se | p-value | 95% CI | | effect |
| alcohol | -0.031 | 0.075 | 0.682 | -0.179 | 0.117 | -0.0209 |
| smoker | 0.109 | 0.088 | 0.214 | -0.063 | 0.282 | 0.082 |
| bmi | 0.705 | 0.783 | 0.368 | -0.832 | 2.24 | 0.617 |
| education° | 0.036 | 0.082 | 0.664 | -0.125 | 0.196 | 0.027 |

*covariates: age, sex, blood processing day, rack barcode
**covariates: age, sex, blood processing day, rack barcode, other self-reported phenotypes (e.[illegible]
°For these models, education was treated as the exposure and shift work was the outcome; e[illegible]

ng Society

| | Model 2 (n=1,032)** | | | |
|---|---|---|---|---|
| se | | p-value | 95% CI | |
| | 0.129 | 0.427 | -0.151 | 0.357 |
| | 0.161 | 0.548 | -0.412 | 0.219 |
| | 1.353 | 0.159 | -0.747 | 4.56 |
| | 0.179 | 0.823 | -0.312 | 0.392 |

| | Model 2 (n=1,032)** | | | |
|---|---|---|---|---|
| se | | p-value | 95% CI | |
| | 0.067 | 0.896 | -0.122 | 0.140 |
| | 0.083 | 0.612 | -0.121 | 0.205 |
| | 0.699 | 0.178 | -0.429 | 2.313 |
| | 0.081 | 0.996 | -0.012 | 0.010 |

| | Model 2** | | | |
|---|---|---|---|---|
| se | | p-value | 95% CI | |
| | 0.076 | 0.785 | -0.171 | 0.129 |
| | 0.094 | 0.382 | -0.102 | 0.267 |
| | 0.797 | 0.439 | -0.947 | 2.181 |
| | 0.092 | 0.77 | -0.154 | 0.208 |

;. for smoking, model adjusted for alcohol, body mass index and education)
ffect estimates are log odds ratios

Supplementary Table 3: Associations between phenotypes and methylation scores in Generati

| | Set 1 (Wave 1) | | | | | |
|---|---|---|---|---|---|---|
| Phenotype | Model 1* (n=1343-2567) | | | | | |
| | effect | se | p-value | 95% CI | | effect |
| alcohol | 0.648 | 0.065 | 9.36E-23 | 0.521 | 0.775 | 0.674 |
| smoker | 0.939 | 0.017 | <1E-100 | 0.906 | 0.972 | 0.916 |
| bmi | 0.144 | 0.002 | <1E-100 | 0.140 | 0.148 | 0.148 |
| education | 0.241 | 0.011 | 2.44E-87 | 0.219 | 0.263 | 0.201 |

*covariates: age, sex and 20 methylation principal components

**covariates: age, sex,  20 methylation principal components, other self-reported phenotypes on Scotland

| Model 2** (n=936-943) | | | | Model 1* (n=2533-[illegible] | | |
|---|---|---|---|---|---|---|
| se | p-value | 95% CI | | effect | se | p-value |
| 0.079 | 6.52E-17 | 0.519 | 0.829 | 0.273 | 0.048 | 1.65E-08 |
| 0.029 | <1E-100 | 0.859 | 0.973 | 0.898 | 0.013 | <1E-100 |
| 0.004 | <1E-100 | 0.140 | 0.156 | 0.148 | 0.004 | <1E-100 |
| 0.013 | 2.74E-45 | 0.176 | 0.226 | 0.201 | 0.013 | 2.36E-25 |

(e.g. for smoking-related DNAm scores, model adjusted for alcohol, body mass index and educ

| Set 2 (Wave 3) | | | | | | |
|---|---|---|---|---|---|---|
| 4423) | | Model 2** (n=2432-2415) | | | | |
| 95% CI | | effect | se | p-value | 95% CI | |
| 0.179 | 0.367 | 0.279 | 0.049 | 2.36E-08 | 0.183 | 0.375 |
| 0.873 | 0.923 | 0.842 | 0.019 | <1E-100 | 0.805 | 0.879 |
| 0.140 | 0.156 | 0.093 | 0.004 | <1E-100 | 0.085 | 0.101 |
| 0.176 | 0.226 | 0.059 | 0.009 | 3.29E-09 | 0.041 | 0.077 |

ation)

Supplementary Table 4: Associations between phenotypes and methylation scores in Understa

| Phenotype | Model 1 (n=1,086-1,167)* | | | | | |
|---|---|---|---|---|---|---|
| | effect | se | p-value | 95% CI | | effect |
| alcohol | 0.321 | 0.047 | 1.29E-11 | 0.228 | 0.414 | 0.324 |
| smoker | 0.908 | 0.031 | <1E-100 | 0.849 | 0.969 | 0.895 |
| bmi | 0.07 | 0.004 | 1.16E-59 | 0.062 | 0.079 | 0.07 |
| education | 0.071 | 0.017 | 3.18E-05 | 0.037 | 0.104 | 0.057 |

*covariates: age, sex, blood processing day, rack barcode

**covariates: age, sex, blood processing day, rack barcode, other self-reported phenotypes (e.[illegible] ınding Society

| | Model 2 (n=1032)** | | | |
|---|---|---|---|---|
| se | | p-value | 95% CI | |
| | 0.049 | 2.74E-11 | 0.228 | 0.419 |
| | 0.033 | <1E-100 | 0.829 | 0.961 |
| | 0.005 | 3.30E-50 | 0.061 | 0.079 |
| | 0.008 | 0.002 | 0.022 | 0.094 |

ʒ. for smoking-related DNAm scores, model adjusted for alcohol, body mass index and educatio ɔn)

Supplementary Table 5: Associations between night shift work and methylation scores in Generatic

| Shiftwork | Methylation score | Model 1* (n=1139-1564) | | | |
|---|---|---|---|---|---|
| | | effect | se | p-value | 95% CI |
| Night hours | alcohol | 0.003 | 0.011 | 0.756 | -0.018 |
| | smoker | 0.040 | 0.010 | 1.28E-04 | 0.019 |
| | bmi | 0.029 | 0.011 | 0.011 | 0.007 |
| | education° | -0.207 | 0.063 | 0.001 | -0.330 |
| Binary night shift work | alcohol | -0.006 | 0.084 | 0.947 | -0.170 |
| | smoker | 0.233 | 0.080 | 0.004 | 0.075 |
| | bmi | 0.219 | 0.089 | 0.014 | 0.044 |
| | education°° | -0.366 | 0.124 | 0.003 | -0.609 |

*covariates: age, sex and 20 methylation principal components

**covariates: age, sex,  20 methylation principal components, other self-reported phenotypes (e.g.

***covariates: age, sex,  20 methylation principal components, corresponding and other self-report

°For these models, education was treated as the exposure and shift work was the outcome; ° effec

›n Scotland

| | Set 1 (Wave 1) | | | | | | |
|---|---|---|---|---|---|---|---|
| | Model 2** (n=481-1078) | | | | | | |
| | effect | se | p-value | 95% CI | | | effect |
| 0.025 | -0.015 | 0.014 | 0.286 | -0.042 | 0.012 | | -0.009 |
| 0.060 | 0.014 | 0.017 | 0.429 | -0.019 | 0.047 | | 0.006 |
| 0.051 | -0.007 | 0.019 | 0.734 | -0.044 | 0.030 | | -0.009 |
| -0.084 | -0.425 | 0.100 | 2.34E-05 | -0.621 | -0.229 | | -0.394 |
| 0.159 | -0.156 | 0.110 | 0.158 | -0.372 | 0.060 | | -0.014 |
| 0.390 | 0.103 | 0.136 | 0.452 | -0.164 | 0.370 | | 0.049 |
| 0.394 | -0.107 | 0.155 | 0.491 | -0.411 | 0.197 | | -0.093 |
| -0.123 | -0.894 | 0.222 | 5.63E-05 | -1.329 | -0.459 | | -0.628 |

for smoking-related DNAm scores, model adjusted for alcohol, body mass index and education
ed phenotypes (e.g. for smoking-related DNAm scores, model adjusted for smoking, alcohol, b
t estimates are log odds ratios

| Model 3*** (n=476-610) | | | | Model 1* (n=1785- | | |
|---|---|---|---|---|---|---|
| se | p-value | 95% CI | | effect | se | p-value |
| 0.016 | 0.606 | -0.040 | 0.022 | -0.003 | 0.008 | 0.708 |
| 0.011 | 0.599 | -0.016 | 0.028 | 0.021 | 0.008 | 0.013 |
| 0.012 | 0.410 | -0.033 | 0.015 | 0.025 | 0.009 | 0.004 |
| 0.140 | 0.005 | -0.668 | -0.120 | -0.087 | 0.050 | 0.082 |
| 0.129 | 0.296 | -0.267 | 0.239 | -0.045 | 0.064 | 0.483 |
| 0.091 | 0.595 | -0.129 | 0.227 | 0.117 | 0.065 | 0.073 |
| 0.097 | 0.339 | -0.283 | 0.097 | 0.209 | 0.069 | 0.002 |
| 0.321 | 0.050 | -1.257 | 0.001 | -0.144 | 0.088 | 0.102 |

)

ody mass index and education)

| | | Set 2 (Wave 3) | | | | |
|---|---|---|---|---|---|---|
| 2428) | | Model 2** (n=1112-2319) | | | | |
| 95% CI | | effect | se | p-value | 95% CI | |
| -0.019 | 0.013 | -0.003 | 0.008 | 0.698 | -0.019 | 0.013 |
| 0.004 | 0.037 | 0.028 | 0.011 | 0.011 | 0.007 | 0.049 |
| 0.008 | 0.043 | 0.014 | 0.011 | 0.210 | -0.007 | 0.036 |
| -0.185 | 0.011 | -0.126 | 0.073 | 0.082 | -0.269 | 0.016 |
| -0.172 | 0.081 | -0.043 | 0.067 | 0.517 | -0.174 | 0.088 |
| -0.011 | 0.245 | 0.157 | 0.085 | 0.063 | -0.010 | 0.324 |
| 0.075 | 0.344 | 0.085 | 0.088 | 0.331 | -0.087 | 0.257 |
| -0.316 | 0.028 | -0.219 | 0.138 | 0.114 | -0.489 | 0.051 |

| Model 3*** (n=1104-1400) | | | | |
|---|---|---|---|---|
| effect | se | p-value | 95% CI | |
| 0.002 | 0.100 | 0.824 | -0.194 | 0.198 |
| 0.012 | 0.007 | 0.095 | -0.002 | 0.026 |
| 0.006 | 0.010 | 0.534 | -0.014 | 0.026 |
| -0.062 | 0.082 | 0.451 | -0.222 | 0.099 |
| -0.014 | 0.079 | 0.863 | -0.169 | 0.141 |
| 0.062 | 0.057 | 0.277 | -0.050 | 0.174 |
| 0.021 | 0.079 | 0.788 | -0.134 | 0.176 |
| -0.109 | 0.161 | 0.501 | -0.425 | 0.207 |

Supplementary Table 6: Associations between night shift work and methylation scores in Understandin

| Methylation scores | Model 1 (n=1,171)* | | | | | |
|---|---|---|---|---|---|---|
| | effect | se | p-value | 95% CI | | effect |
| alcohol | -0.029 | 0.201 | 0.884 | -0.432 | 0.365 | 0.011 |
| smoker | 0.063 | 0.218 | 0.771 | -0.364 | 0.492 | 0.029 |
| bmi | 0.008 | 0.222 | 0.97 | -0.427 | 0.443 | 0.038 |
| education° | -0.105 | 0.265 | 0.691 | -0.624 | 0.413 | -0.102 |

| Methylation scores | Model 1 (n=1,171)* | | | | | |
|---|---|---|---|---|---|---|
| | effect | se | p-value | 95% CI | | effect |
| alcohol | -0.021 | 0.099 | 0.831 | -0.216 | 0.174 | -0.053 |
| smoker | 0.136 | 0.108 | 0.208 | -0.076 | 0.348 | 0.154 |
| bmi | 0.073 | 0.109 | 0.503 | -0.142 | 0.289 | 0.087 |
| education° | 0.028 | 0.127 | 0.826 | -0.22 | 0.276 | 0.052 |

| Methylation scores | Model 1 (n=1,171)* | | | | | |
|---|---|---|---|---|---|---|
| | effect | se | p-value | 95% CI | | effect |
| alcohol | -0.017 | 0.111 | 0.879 | -0.235 | 0.201 | -0.068 |
| smoker | 0.149 | 0.119 | 0.213 | -0.086 | 0.384 | 0.184 |
| bmi | 0.088 | 0.123 | 0.473 | -0.153 | 0.329 | 0.1 |
| education° | 0.069 | 0.141 | 0.627 | -0.208 | 0.345 | 0.103 |

*covariates: age, sex, blood processing day, rack barcode
**covariates: age, sex, blood processing day, rack barcode, other self-reported phenotypes (e.g. for smo
***covariates: age, sex, blood processing day, rack barcode, corresponding other self-reported phenoty
°For these models, education was treated as the exposure and shift work was the outcome; effect esti

g Society

**CURRENT NIGHT SHIFT WORK**

| Model 2 (n=1,032-1,107)** | | | | | | Model 3 (n=1,032) |
|---|---|---|---|---|---|---|
| se | p-value | 95% CI | | effect | se | p-value |
| 0.199 | 0.597 | -0.379 | 0.402 | -0.03 | 0.2 | 0.755 |
| 0.223 | 0.182 | -0.408 | 0.467 | 0.116 | 0.17 | 0.188 |
| 0.225 | 0.446 | -0.404 | 0.479 | -0.087 | 0.203 | 0.902 |
| 0.296 | 0.73 | -0.683 | 0.478 | -0.356 | 0.297 | 0.904 |

**EVER NIGHT SHIFT WORK**

| Model 2 (n=1,032-1,107)** | | | | | | Model 3 (n=1,032) |
|---|---|---|---|---|---|---|
| se | p-value | 95% CI | | effect | se | p-value |
| 0.11 | 0.954 | -0.25 | 0.144 | -0.032 | 0.103 | 0.88 |
| 0.115 | 0.894 | -0.072 | 0.379 | 0.116 | 0.088 | 0.496 |
| 0.114 | 0.867 | -0.137 | 0.312 | 0.013 | 0.105 | 0.669 |
| 0.142 | 0.715 | -0.226 | 0.33 | 0.253 | 0.258 | 0.326 |

**PREVIOUS NIGHT SHIFT WORK**

| Model 2 (n=1,032-1,107)** | | | | | | Model 3 (n=1,032) |
|---|---|---|---|---|---|---|
| se | p-value | 95% CI | | effect | se | p-value |
| 0.112 | 0.547 | -0.289 | 0.153 | 0.012 | 0.116 | 0.918 |
| 0.129 | 0.156 | -0.071 | 0.438 | 0.16 | 0.099 | 0.106 |
| 0.129 | 0.844 | -0.154 | 0.355 | 0.044 | 0.119 | 0.713 |
| 0.161 | 0.521 | -0.212 | 0.418 | 0.091 | 0.162 | 0.575 |

oking DNAm, model adjusted for alcohol, body mass index and education)
pes (e.g. for smoking DNAm, model adjusted for smoking, alcohol, body mass index and educa
mates are log odds ratios

| )*** | |
|---|---|
| 95% CI | |
| -0.423 | 0.363 |
| -0.218 | 0.451 |
| -0.487 | 0.313 |
| -0.617 | 0.546 |

| )*** | |
|---|---|
| 95% CI | |
| -0.235 | 0.171 |
| -0.056 | 0.289 |
| -0.193 | 0.219 |
| -0.252 | 0.758 |

| )*** | |
|---|---|
| 95% CI | |
| -0.215 | 0.239 |
| -0.034 | 0.354 |
| -0.191 | 0.279 |
| -0.227 | 0.409 |

tion)

Supplementary Table 7: Associations between night shift work and measures of epigenetic age accele

| Shift work | Clock | Set 1 | | | |
|---|---|---|---|---|---|
| | | Model 1* (n=1142-1564) | | | Mode |
| | | effect | se | p-value | effect |
| **Night hours** | **Horvath** | -0.027 | 0.04 | 0.503 | -0.061 |
| | **IEAA** | -0.03 | 0.04 | 0.453 | -0.065 |
| | **Hannum** | -0.021 | 0.031 | 0.486 | -0.02 |
| | **EEAA** | -0.021 | 0.041 | 0.603 | -0.013 |
| | **PhenoAge** | 0.058 | 0.046 | 0.211 | -0.002 |
| | **GrimAge** | 0.19 | 0.043 | 1.18E-05 | 0.119 |
| **Binary night shift work** | **Horvath** | -0.033 | 0.314 | 0.916 | -0.423 |
| | **IEAA** | -0.030 | 0.314 | 0.924 | -0.417 |
| | **Hannum** | -0.277 | 0.243 | 0.255 | -0.502 |
| | **EEAA** | -0.383 | 0.320 | 0.231 | -0.625 |
| | **PhenoAge** | 0.356 | 0.362 | 0.325 | -0.489 |
| | **GrimAge** | 0.983 | 0.340 | 0.004 | 0.661 |

*covariates: sex and 20 methylation principal components

**covariates: sex, 20 methylation PCs, alcohol, bmi, education, smoker

IEAA: Instrinsic epigenetic age acceleration

EEAA: Extrinsic epigenetic age acceleration

ration in *Generation Scotland* dataset

| | | **Set 2** | | | | |
|---|---|---|---|---|---|---|
| **el 2** (n=739-1025)** | | **Model 1* (n=1785-2428)** | | | **Model 2** (n=1623-** | |
| se | p-value | effect | se | p-value | effect | se |
| 0.053 | 0.251 | -0.009 | 0.031 | 0.758 | -0.013 | 0.031 |
| 0.053 | 0.215 | -0.014 | 0.031 | 0.621 | -0.017 | 0.031 |
| 0.039 | 0.605 | 0.012 | 0.027 | 0.631 | 0.009 | 0.027 |
| 0.051 | 0.807 | 0.035 | 0.035 | 0.289 | 0.03 | 0.035 |
| 0.059 | 0.969 | 0.05 | 0.043 | 0.206 | 0.061 | 0.043 |
| 0.04 | 3.13E-03 | 0.101 | 0.024 | 9.45E-04 | 0.052 | 0.024 |
| 0.424 | 0.318 | 0.043 | 0.221 | 0.848 | 0.010 | 0.241 |
| 0.423 | 0.324 | -0.017 | 0.220 | 0.940 | -0.035 | 0.241 |
| 0.315 | 0.111 | 0.206 | 0.192 | 0.282 | 0.174 | 0.206 |
| 0.408 | 0.126 | 0.395 | 0.255 | 0.121 | 0.352 | 0.275 |
| 0.464 | 0.292 | 0.432 | 0.315 | 0.170 | 0.534 | 0.340 |
| 0.331 | 0.046 | 0.713 | 0.241 | 0.003 | 0.348 | 0.186 |

| -2217) |
|---|
| p-value |
| 0.672 |
| 0.592 |
| 0.739 |
| 0.401 |
| 0.15 |
| 0.031 |
| 0.967 |
| 0.886 |
| 0.397 |
| 0.200 |
| 0.117 |
| 0.061 |

Supplementary Table 8: Associations between night shift work and measures of epigenetic ag

| | Current night shift | | | | | |
|---|---|---|---|---|---|---|
| **Clock** | **Model 1* (n = 1,128)** | | | **Model 2** (n = 998)** | | |
| | **beta** | **se** | **p-value** | **beta** | **se** | **p-value** |
| Horvath | 1.402 | 1.026 | 0.172 | 1.386 | 1.043 | 0.184 |
| IEAA | 1.043 | 1.018 | 0.306 | 0.908 | 1.036 | 0.381 |
| Hannum | 0.212 | 0.921 | 0.818 | 0.097 | 0.950 | 0.919 |
| EEAA | 0.016 | 0.875 | 0.985 | -0.161 | 0.902 | 0.859 |
| PhenoAge | 2.420 | 1.345 | 0.072 | 2.174 | 1.335 | 0.104 |
| GrimAge | 0.581 | 1.149 | 0.613 | 0.206 | 1.186 | 0.862 |

*covariates: sex, blood processing day, rack barcode
**covariates: age, sex, blood processing day, rack barcode, alcohol, bmi, education, smoker

IEAA: Instrinsic epigenetic age acceleration
EEAA: Extrinsic epigenetic age acceleration

e acceleration in Understanding Society

| Ever night shift | | | | | | |
|---|---|---|---|---|---|---|
| Model 1* (n = 1,171) | | | Model 2** (n = 1,032) | | | Mo |
| beta | se | p-value | beta | se | p-value | beta |
| 0.223 | 0.446 | 0.618 | 0.305 | 0.473 | 0.520 | -0.013 |
| 0.150 | 0.443 | 0.735 | 0.168 | 0.470 | 0.721 | -0.055 |
| 0.212 | 0.400 | 0.596 | 0.254 | 0.429 | 0.554 | 0.176 |
| 0.013 | 0.380 | 0.973 | 0.079 | 0.408 | 0.847 | -0.067 |
| 0.484 | 0.588 | 0.411 | 0.524 | 0.610 | 0.391 | -0.251 |
| -0.295 | 0.498 | 0.553 | -0.111 | 0.536 | 0.836 | -0.377 |

| Previous night shift | | | | |
|---|---|---|---|---|
| del 1* (n = 1,147) | | Model 2** (n = 1,010) | | |
| se | p-value | beta | se | p-value |
| 0.505 | 0.980 | 0.130 | 0.540 | 0.810 |
| 0.503 | 0.913 | 0.019 | 0.537 | 0.971 |
| 0.452 | 0.696 | 0.283 | 0.488 | 0.563 |
| 0.430 | 0.876 | 0.056 | 0.465 | 0.905 |
| 0.663 | 0.705 | -0.152 | 0.693 | 0.827 |
| 0.563 | 0.503 | -0.040 | 0.609 | 0.948 |
