## Supplementary Methods for "Epigenetic markers of adverse lifestyle identified among night shift workers"

**DNA methylation data in Generation Scotland**

GS comprises >24,000 participants (59% female, average age = 47.2 years (SD = 15.1)) from ~7,000 families who were recruited from the registers of collaborating general practitioners in Scotland between 2006 and 2011. Data were collected on various cognitive, psychiatric and health measures, and blood DNA samples were collected for genotyping and methylation profiling. All components of GS received ethical approval from the NHS Tayside Committee on Medical Research Ethics (REC Reference Number: 05/S1401/89). GS has also been granted Research Tissue Bank status by the East of Scotland Research Ethics Service (REC Reference Number: 20-ES-0021), providing generic ethical approval for a wide range of uses within medical research.

Details of the sample sets, including quality control and normalisation of the DNAm data in each sample, have been described in detail previously^1-3^ The two data sets had DNAm profiled at separate time points and quality control and normalisation of each set was carried out separately. Methylation β values at each CpG site were transformed to obtain *M*-values [log2(β/ (1- β)] for statistical analysis. The final dataset for Set 1 comprised M-values at 777,193 CpGs measured in 5,087 related individuals, while Set 2 comprised M-values at 773,860 CpGs measured in 4,450 individuals, unrelated to each other, and to those in Set 1. To remove the confounding effects of family structure, Set 1 was further filtered to a subset of unrelated individuals (n=2,578).

**DNA methylation data in Understanding Society**

The British Household Panel Survey (BHPS) comprises a clustered random sample of households recruited in 1991 with all members followed annually. In 2010 the BHPS was incorporated into the larger UK Household Longitudinal Study (UKHLS) (also known as *Understanding Society)*.^4,5^ Since 1991, annual interviews have collected sociodemographic information and in 2011-2012, biomedical measures and blood samples for BHPS participants were collected at a nurse visit in the participants’ homes. Respondents were eligible to give a blood sample if they had taken part in the previous main interview in England, were aged 16+ years, lived in England, Wales or Scotland, were not pregnant, and met other conditions detailed in the user guide ^6,7^. DNA was extracted from non-fasting blood samples and stored for genetic and epigenetic analyses.

Details of the DNAm data, including quality control and normalisation, have been described in detail previously.^8^ Methylation β values at each CpG site were transformed to obtain *M*-values [log2(β/ (1- β)] for statistical analysis.

**Lifestyle factors in Generation Scotland**

Information on night shift work was obtained from a question in the baseline questionnaire which asked, “If you are employed, how many hours in a typical week would you work in the evening or overnight?”. An 11-level ordinal variable for night hours was derived from 0 hours to >50 hours evening/overnight work per week. Individuals were excluded if they reported being a housewife/homemaker, retired, a full-time student, unemployed, other, or not applicable for their job status. We derived a binary variable for “night shift work” based on ≥20 hours/week compared with those reporting to work 0 hours in the evening or overnight (i.e., excluding 1-19 hours of night shift work).

Information on alcohol intake, smoking and education were taken from a questionnaire administered at baseline and prior to a clinic visit, where BMI (the ratio of weight in kilograms to height in metres squared (kg/m^2^)) was measured and blood for DNA extraction was taken.

Participants were asked on average how often do they drink alcohol. Those who responded: ‘daily or almost daily’ were classed as “daily” drinkers; ‘3 or 4 days per week’ or ‘1 or 2 days per week’ as “more than weekly” drinkers; ‘1-3 days per month’ as “more than monthly” drinkers; ‘special occasions only’ or ‘rarely’ were classed as “less than monthly” drinkers.

Participants were asked whether they had ever smoked tobacco. Those who responded, ‘yes, currently smoke’, were classed as “current smokers”; ‘yes, but stopped within past 12 months’ or ‘yes, but stopped more than 12 months ago’ as “former smokers”; ‘no, never smoked’ as “never smokers”.

Participants were asked how many years they attended school or studied full-time. Participants chose from 0 years; 1-4 years; 5-9 years; 10-11 years; 12-13 years; 14-15 years; 16-17 years; 18-19 years; 20-21 years; 22-23 years or >24 years of full-time education. The midpoints of categories were then calculated.

**Lifestyle factors in Understanding Society**

Repeated measures of reported night shift work was available in UKHLS, unlike in GS where this data was only provided at one time point. Questionnaire data were obtained from annual surveys conducted as part of BHPS (waves 9-18; 1999-2009) and UKHLS (wave 2; 2010). At waves 9-12 and 14-18 of BHPS, and wave 2 of UKHLS, participants were asked in the Employment section of the survey which category best described the times of day usually worked: “Mornings only”, “Afternoons only”, “During the day”, “Evenings only”, “At night”, “Both lunchtimes and evening”, “Other times of day”, “Rotating shifts”, “Varies/no usual pattern”, “Daytimes and evenings”, “Other (please give details)”. Those individuals who reported that they worked evenings only or at night were classed as “night shift workers”, while those reporting other times of work were classed as “non-night shift workers”. We used data at 10 time points to derive three measures of shift work: ever worked night shifts, currently working night shifts (at wave 2 of UKHLS) and previously worked night shifts.

Information on alcohol intake, smoking and education was taken from questionnaire data and BMI was determined at the nurse visit during waves 2 and 3.

In wave 2 of UKHLS, participants were asked how often they had an alcoholic drink during the last 12 months. Those who reported drinking: ‘almost every day’ or ‘five or six days a week’ were classed as “daily” drinkers; ‘three or four days a week’ or ‘once or twice a week’ as “more than weekly” drinkers; ‘once or twice a month’ as “more than monthly” drinkers; ‘once every couple of months’, ‘once or twice a year’, ‘not at all in the last 12 months’ or ‘not in the last 12 months’ as “less than monthly” drinkers.

Participants were asked about their smoking history and whether they had ever smoked a cigarette, a cigar, or a pipe. Participants who responded with ‘no’ were classed as “never smokers”; ‘yes’ and that they were currently smoking as “current smokers”; ‘yes’ but not currently smoking as classed as “former smokers”.

Participants were asked what their current highest educational qualification was. Years of education was assigned based on the highest educational qualification achieved, where ‘degree’, ‘other higher degree’, ‘A-level etc.’, ‘GCSE etc.’ and ‘other qualification’ were assigned 20, 19, 13, 10 and 7 years of education, respectively.

**Epigenetic age measures**

To generate the epigenetic ageing measures, we used the DNA Methylation Age Calculator (<https://dnamage.genetics.ucla.edu/>) developed by the Horvath lab. We obtained six estimates of DNAm age as well as measures of age acceleration, i.e., the deviation of predicted DNAm age from chronological age, based on the six predictors. The six epigenetic age measures derived in GS and UKHLS were: Horvath’s Epigenetic Age Acceleration (AgeAccelHorvath); Horvath’s multi-tissue predictor of Intrinsic Epigenetic Age Acceleration (IEAA), Hannum’s predictor of Epigenetic Age Acceleration (AgeAccelHannum); Extrinsic Epigenetic Age Acceleration (EEAA), an enhanced version of the Hannum predictor which up-weights the contribution of blood cell composition; PhenoAge Acceleration (AgeAccelPheno), optimised to predict physiological dysregulation; and GrimAge Acceleration (AgeAccelGrim), optimised to predict lifespan.

6.	Benzeval M, Davillas A, Kumari M, Lynn P. *Understanding Society: the UK Household Longitudinal Study biomarker user guide and glossary.* Institute for Social and Economic Research, University of Essex 2014.

7.	University of Essex. Institute for Social and Economic Research NSR. Understanding Society: Waves 2 and 3 Nurse Health Assessment, 2010- 2012 [data collection].3rd Edition. UK Data Service. SN:7251	<http://doi.org/10.5255/UKDA-SN-7251-3>. In:2014.

8.	Gorrie-Stone TJ, Saffari, A., Malki, K, Schalkwyk, L.C. Bigmelon: Illumina methylation array analysis for large experiment R package version 160. 2018; <https://rdrr.io/bioc/bigmelon/f/inst/doc/bigmelon.pdf>, 04/06/19.
